# Three Mutations Convert the Selectivity of a Protein Sensor from Nicotinic Agonists to S-methadone for Use in Cells, Organelles, and Biofluids

**DOI:** 10.1101/2022.02.24.481226

**Authors:** Anand K. Muthusamy, Charlene H. Kim, Scott C. Virgil, Hailey J. Knox, Jonathan S. Marvin, Aaron L. Nichols, Bruce N. Cohen, Dennis A. Dougherty, Loren L. Looger, Henry A. Lester

## Abstract

We report a reagentless, intensity-based S-methadone fluorescent sensor, iS-methadoneSnFR, consisting of a circularly permuted GFP inserted within the sequence of a mutated bacterial periplasmic binding protein (PBP). We evolved a previously reported nicotine-binding PBP to become a selective S-methadone-binding sensor, via three mutations in the PBP’s second shell and hinge regions. iS-methadoneSnFR displays the necessary sensitivity, kinetics, and selectivity – notably enantioselectivity against R-methadone – for biological applications. Robust iS-methadoneSnFR responses in human sweat and saliva and mouse serum enable diagnostic uses. Expression and imaging in mammalian cells demonstrate that S-methadone enters at least two organelles and undergoes acid trapping in the Golgi apparatus, where opioid receptors can signal. This work shows a straightforward strategy in adapting existing PBPs to serve real-time applications ranging from subcellular to personal pharmacokinetics.

---

We report the first selective real-time fluorescent biosensor for a small molecule opioid, “intensity-based <u>S-methadone s</u>e<u>n</u>sing fluorescent reporter” or “iS-methadoneSnFR” (**Figure 1**). To employ the indicator for quantitative dynamic opioid measurements in cells and biofluids, we engineered iS-methadoneSnFR to meet necessary criteria: (1) sensitivity in the pharmacological range, (2) selectivity against endogenous molecules, (3) selectivity against exogenous drugs, including those of the same drug class, (4) photostability for the duration of measurements, (5) physical stability outside cells, and (6) reversible binding with ∼second resolution.

**Figure 1:**
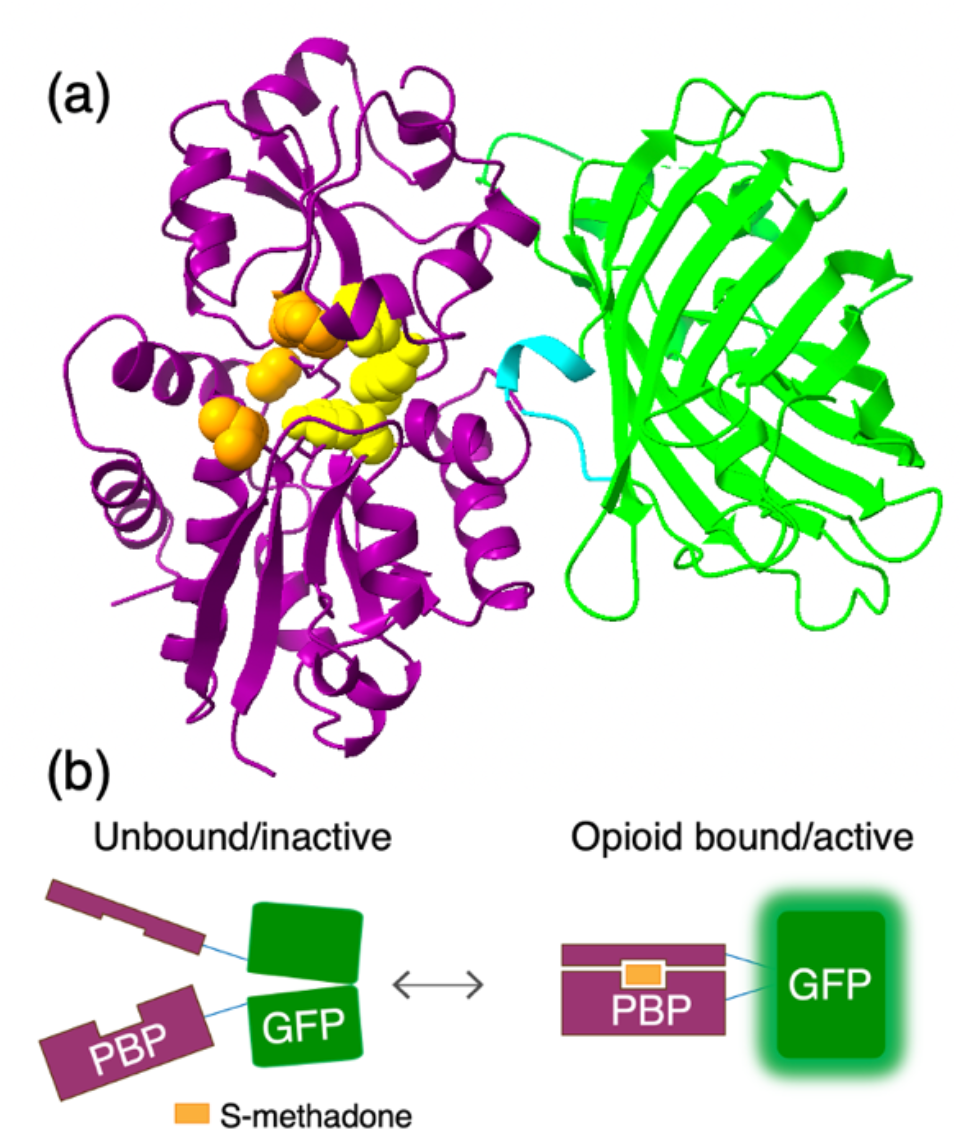
(a) Crystal structure of iNicSnFR3a (PDB: 7S7T) mutated *in silico* to iS-methadoneSnFR (mutations shown in orange spheres). All but one putative cation-π residue in iNicSnFR3a were maintained in iS-methadoneSnFR’s binding pocket (critical residues Y65, Y357, and Y460 shown as yellow spheres). (b) Biosensor mechanism: in the unbound state, GFP’s chromophore has a poor environment for fluorescence. The PBP binds S-methadone with a “Venus flytrap” conformational change, increasing the brightness of the GFP chromophore.

The risk of opioid-use disorder and death by overdose has increased alongside the worldwide access to highly potent opioid agonists^1^. Nevertheless, opioids remain essential analgesics. Since the 1960s, methadone maintenance therapy (MMT) has served to reduce harm from opioid addiction^2,3^. MMT relies on pharmacokinetics: oral methadone’s onset is slower than that of injected or inhaled μ-opioids, and its effects last much longer due to a ∼24 h half-life^4^. Therefore, despite acting as a μ-opioid agonist, methadone staves off withdrawal symptoms without producing the euphoria associated with other agonists^4^. However, interindividual variability in the metabolism of methadone, partially due to polymorphisms in cytochrome P450 isotypes^5,6^, can lead to therapeutic failures^7^. Methadone is clinically administered as the racemate and measuring either enantiomer is suitable for therapeutic drug monitoring^8^. Drug metabolism is conventionally addressed by blood draw, but this method is laborious, invasive, and restricted to the clinic. An optimal methadone readout would enable personalized dosing regimens, by producing real-time tracking of [methadone] in biological fluids and facilitating tapering from potent opioids. Within a subject, opioid pharmacokinetics also varies at the level of intracellular compartments to produce acid trapping and diverse interactions with receptors including chaperoning and activation^9,10,11^. In both cases, a sensor with *in situ* readout and ∼second resolution is required.

Conventional small molecule detection methods have been extended to methadone but may be limited in specificity, time resolution, or spatial resolution^12^. An antibody against methadone was used in a lateral flow test of human sweat (limited to a single timepoint)^13^. Electrochemical methods provide continuous measurements but vary in selectivity against other opioids^14-16^ and, in all cases, cannot be used for subcellular measurements. A pioneering de novo protein design campaign for an opioid sensor, binding of fentanyl produced a conformational switch in a transcription factor^17^ but required a cellular readout and hours-to-days temporal resolution.

We hypothesized that all the required criteria could be satisfied by a single-chain sensor comprising a mutated bacterial periplasmic binding protein (PBP), a variant of the choline-binding protein OpuBC from *Thermoanaerobacter sp513*, interrupted by a circularly permuted GFP (cpGFP)(**Figure 1**)^18-20^. The cpGFP insertion approach has also been used in Ca^2+^ sensors (the GCaMP series) and in neurotransmitter sensors^18,21^. Our strategy consisted of (1) screening each methadone enantiomer against a previously reported nicotine biosensor, iNicSnFR3a, and its variants^19^ and (2) iterative site-saturation mutagenesis to select for S-methadone and against cholinergic ligands (**Figure 2a**). We performed chiral resolution on racemic methadone to isolate (+)-S-methadone and (-)-R-methadone (assigned by optical rotation^22^) with analytical purity and 99% enantiomeric excess (**Figure S1**).

**Figure 2:**
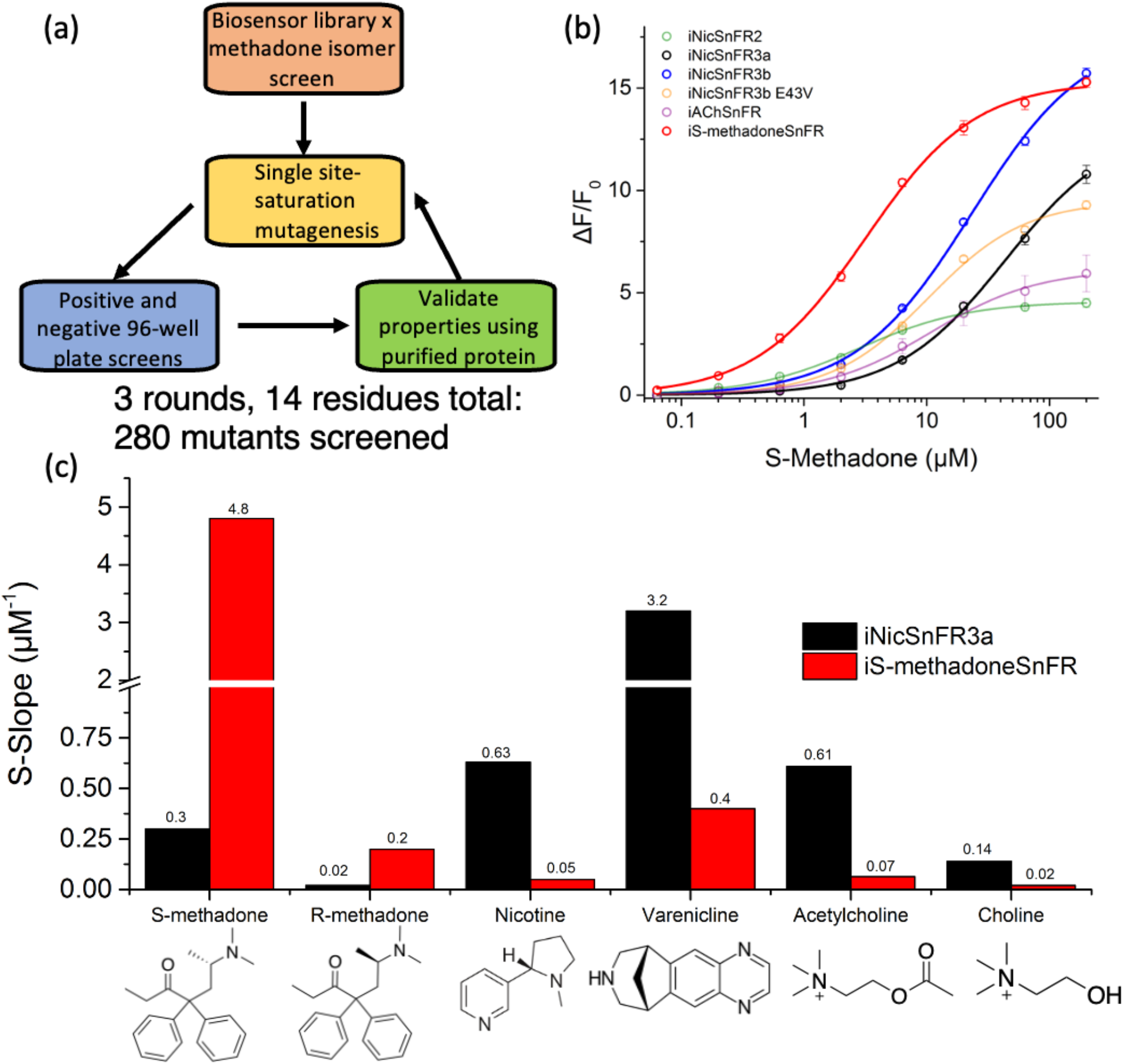
(a) Directed evolution strategy. (b) Fluorescence responses to S-methadone. iNicSnFR3a (black) has several variants (faded curves), of which one has markedly better sensitivity, owing to the N11E mutation (blue). This lead was evolved to iS-methadoneSnFR (red), which included re-optimization at position 11. Only the final biosensor had sufficient sensitivity at 1 μM (vertical black line; the relevant maintenance concentration). (c) Shift in selectivity from iNicSnFR3a (black) to iS-methadoneSnFR (red) measured by S-slope (see text). Note the scale change at the axis break.

While there is no structural homology or pharmacological overlap between nicotinic and μ-opioid receptors, several variants of nicotinic drug biosensors displayed weak fluorescence responses to S-methadone (**Figure 2b**). Although the PBP had no enantioselective pressure for binding its achiral ligand choline, all variants screened to date displayed enantioselectivity for S-methadone (**Figure S2**). Dose-response relations were fit to the Hill equation to determine an EC_50_ and ΔF-_max_/F_0_. In the linear portion of the dose-response relation we define the increase in fluorescence per micromolar, “S-slope”, as a metric of biosensor sensitivity: (Δ(F/F_0_)/(Δ[ligand]) at [drug] << EC50^23^). For a Hill coefficient of ∼1.0, the S-slope equals the ratio (ΔF-_max_/F_0_)/EC_50_. A variant of iNicSnFR3a, iNicSnFR3b, provided the largest dynamic range for both S-methadone and R-methadone (**Figure S2**) and served as the input to several rounds of directed evolution.

We selected for both an increase in sensitivity to S-methadone and a decrease in sensitivity to nicotinic ligands. We chose mutation sites based on a crystal structure of iNicSnFR1 (PDB: 6EFR) and directed evolution of iNicSnFR3a14. The resulting sensor displayed a ∼16-fold improvement in sensitivity over iNicSnFR3a; ΔF/F_0_ increased to 3.76 ± 0.16 at 1 μM, the representative plasma maintenance concentration^8^ (**Figure 2b**). Notably, iS-methadoneSnFR displayed sensitivity to S-methadone that exceeded the sensitivity for any of the original cholinergic ligands and displayed a marked shift in ligand selectivity (**Figure 2c**). iS-methadoneSnFR displayed near-zero response for physiologically or pharmacologically relevant steady-state acetylcholine (ACh), choline, varenicline, and nicotine concentrations (∼1 μM, 10 to 20 μM^24^, 0 to 100 nM^25^ and ∼25 to ∼500 nM, respectively^26^ (**Figure 4a** below)).

**Figure 3:**
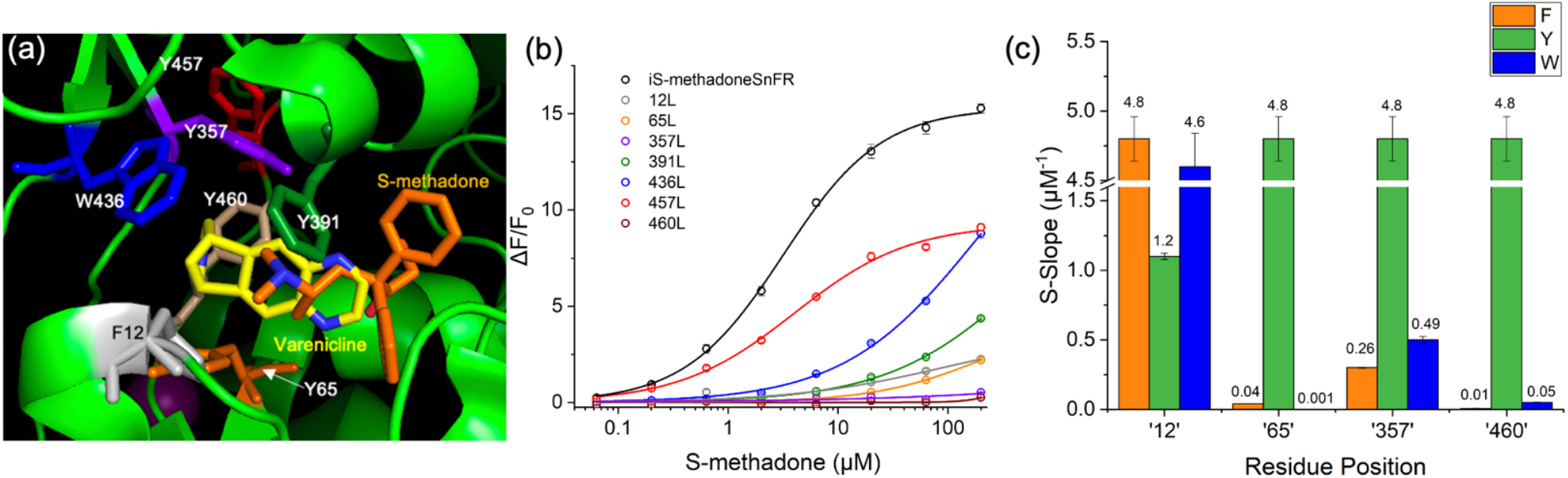
(a) PDB: 7S7T (iNicSnFR3a, varenicline bound) showing cation-π interactions with Y65 and Y357. S-methadone was docked into 7S7T. (b) Fluorescence dose-response relations of cation-π residue Leu mutants. (c) Aromatic side-chain screen through critical positions identified in (b) with resulting S-slope. Note the break in y-axis.

**Figure 4:**
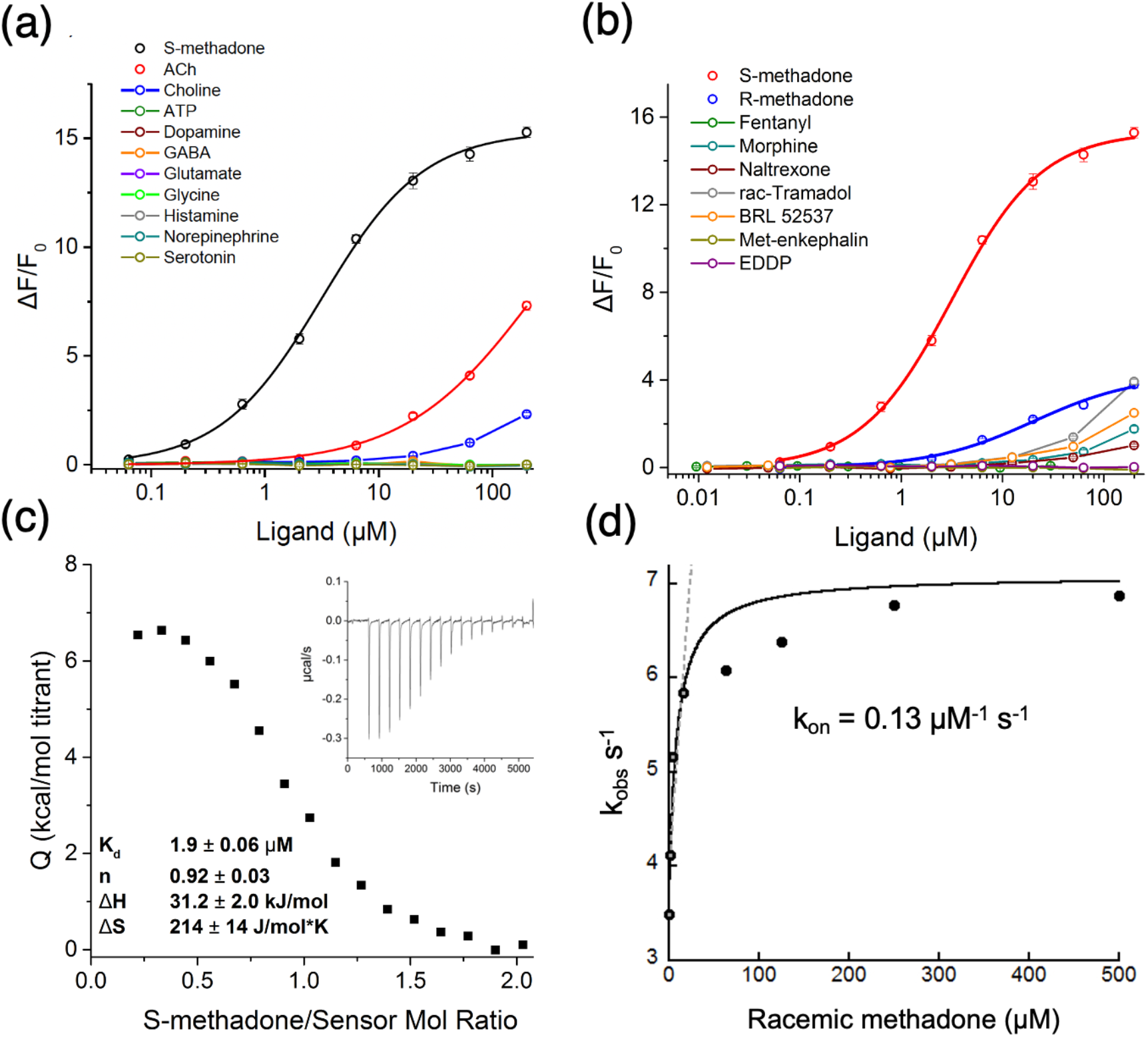
Selectivity and biophysical properties of iS-methadoneSnFR. (a) iS-methadoneSnFR vs endogenous neurotransmitters and choline. Responses to ACh and choline had S-slopes <0.1 μM^-1^. (b) iS-methadoneSnFR vs other clinically used opioids. The response to R-methadone was near zero at ∼1 μM. Weak or no responses were observed for other drugs tested. EDDP is 2-Ethylidene-1,5-dimethyl-3,3-diphenylpyrrolidine, the major metabolite of methadone. (c) Isothermal titration calorimetry of purified iS-methadoneSnFR. 30 μM of the biosensor was mixed with 2 μM injections of 300 μM S-methadone. (d) Stopped-flow kinetic measurements with racemic methadone.

We characterized iS-methadoneSnFR’s binding using docking and biochemical studies. Although only three mutations were required to generate iS-methadoneSnFR from iNicSnFR3a/3b, advantageous mutations were rare: ∼1% of all mutations screened were accepted as improvements. Docking S-methadone into recently reported structures of liganded iNicSnFR3a^27^ showed that the N-methyl groups of S-methadone lie 4.6 and 5.5 Å from the aromatic groups of Y357 and Y65, respectively (slightly greater than the distance from the beta carbons of varenicline to these two groups). In the initial round of mutations, most sites yielded no improvement, except for a W436F mutation spatially near Y65 and Y357 (Figure S3). We previously reported nicotine and varenicline making cation-π interactions with Y65 and Y357 in iNicSnFR3a (PDB: 7S7T, 7S7U, respectively)^27^. Each nicotinic ligand bears a protonated nitrogen lying midway on the axis of the aromatic centroids of Y65 and Y357 (**Figure 3a**). In the subsequent round, second-shell mutation N11V created additional volume next to F12, in the second shell. Finally, the third round yielded L490A, allowing for greater flexibility in the hinging of the PBP.

Leucine mutagenesis among individual binding pocket aromatic residues showed the primacy of Y65, Y357, F12, and Y460 (**Figure 3b**). An aromatic side-chain screen across these four positions revealed a necessity of Tyr in the 1st shell positions Y65, Y357, and Y460 (**Figure 3c**). Substituting a noncanonical side chain, O-methyltyrosine, yielded a near-null biosensor at residue 65 but not at 12 (**Figure S4**). These data suggest that S-methadone’s amine directly interacts with the 1st shell residues, as with nicotinic drugs, and the phenolic -OH is necessary for hydrogen bonding. The three accepted mutations represent a 94 Å^3^ reduction in van der Waals’ volume, comparable to the 132 Å^3^ increase in ligand volume from varenicline to methadone, as though the accepted mutations allowed S-methadone better access to aromatic residues critical to binding both classes of drugs. Therefore, the PBP has an aromatic binding pocket for protonated amines, and other regions of the binding site can be tuned to accommodate the remainder of the ligand’s steric bulk and functional groups.

iS-methadoneSnFR satisfied our sensitivity, selectivity, and biophysical criteria for a useful biosensor. Fluorescence dose-response relations showed an excellent dynamic range, ΔF_max_/F_0_ of 15.3 ± 0.2, and an EC_50_, 3.2 ± 0.2 μM, near the relevant plasma concentrations for maintenance therapy^8^. Isothermal titration calorimetry (ITC) determined a K_d_ of 1.9 ± 0.2 μM, in good agreement with the fluorescence EC_50_ (**Figure 4c**). ITC also demonstrated a single binding site (stoichiometry = 0.92) with an entropically-driven conformational change. iS-methadoneSnFR had little or no response (S-slope < 0.1 μM^-1^) to other neurotransmitters (**Figure 4a**) and other opioids (**Figure 4b**). The S-slope for S-methadone was ∼20x that for R-methadone. When we added R-methadone to S-methadone, fluorescence was modestly elevated at lower [S-methadone], but all responses converged at the ΔF_max_/F_0_ for S-methadone alone (**Figure S5**). 1-second stopped flow kinetics were obtained using racemic methadone (**Figure S4**) and determined an apparent k_on_ of 0.13 μM^-1^s^-1^ (**Figure 4d**). The final 10 ms of the 1 second stopped-flow traces were fitted by a Hill equation with EC_50_ ∼ 8 μM (**Figure S6**) for the racemate, which was approximately double the EC_50_ for S-methadone alone (as expected if the binding strongly favors the S-enantiomer).

Therapeutic use of opioids would be improved by quantitative, real-time, minimally invasive or non-invasive measurements in sweat, saliva, and interstitial fluid^28,29^. The selectivity and high aqueous solubility of iS-methadoneSnFR enable its use in such applications. We tested the biosensor in PBS:biofluid samples and found robust responses in the pharmacologically relevant concentration range (**Figure 5**). iS-methadoneSnFR, like all GFP-based biosensors, displays smaller responses at pH < ∼7 (**Figure S7**). Because biofluids, particularly sweat, have variable and/or acidic pH, 3x PBS pH 7.4 was used to partially buffer a mixture with the biofluid. Still, the response at 1 μM and below in the biofluids provide at least ∼200% dynamic range.

**Figure 5:**
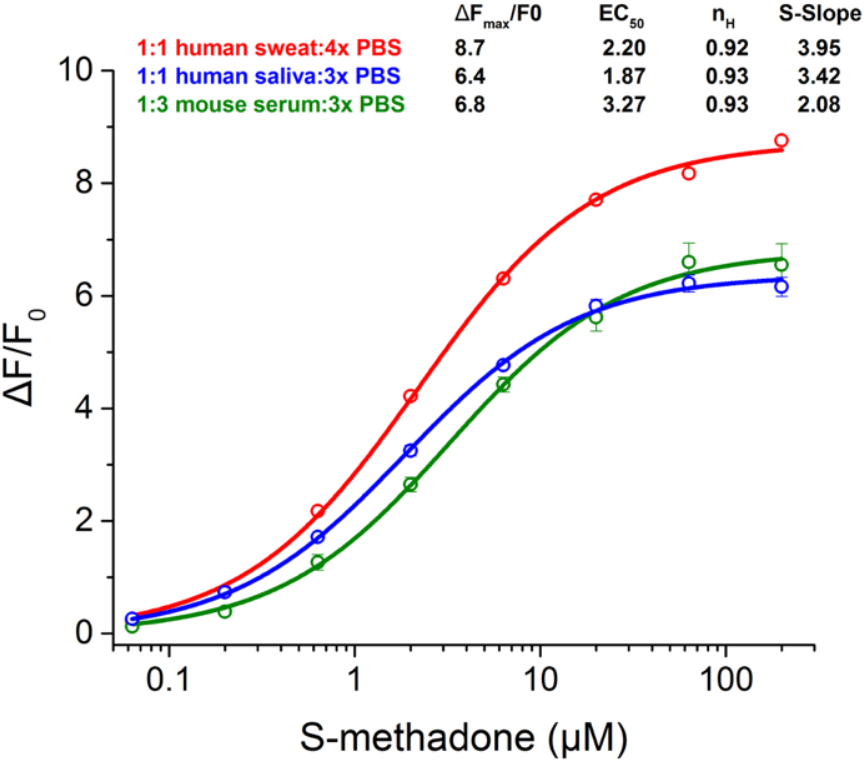
iS-methadoneSnFR dose-response relation in biofluids. 1:1 mixture of drug/biosensor in 3x PBS pH 7.4 with human sweat, human saliva, and 1:3 mixture with mouse serum (no pH adjustment of any biofluid).

At the subcellular level, membrane-permeant weakly basic opioid drugs, but not impermeant derivatives or endogenous opioid peptides, enter the endoplasmic reticulum, and can act as pharmacological chaperones, altering the folding and trafficking of their receptors^10^. Opioid drugs also activate their receptors in endosomes and the Golgi apparatus^11^. We targeted iS-methadoneSnFR to the plasma membrane, endoplasmic reticulum, and Golgi apparatus of HeLa using targeting sequences (**Figure 6a**). We applied pulses of S-methadone (0 to 250 nM in 50 nM steps) to measure the linear portion of the dose-response relation (S-slope) in wide-field imaging (**Figure S8**). The results indicate that ample S-methadone is available in the ER for potential chaperoning. The Golgi showed the largest S-slope among the three compartments (1.7x that of PM), despite having the lowest pH (**Figure 6b**). After correcting the S-slope for pH dependence, we find an accumulation factor of 2.9x to 4.4x across the Golgi pH range of 6.3 to 6.8^25^ (**Figure S8**). Accumulation of opioids such as methadone in acidic compartments^31^ may lead to intensified G-protein coupled signaling. We also validated iS-methadoneSnFR for time-resolved measurements in primary hippocampal neurons, encouraging mechanistic studies in tissues and *in vivo* (**Figure S9**).

**Figure 6:**
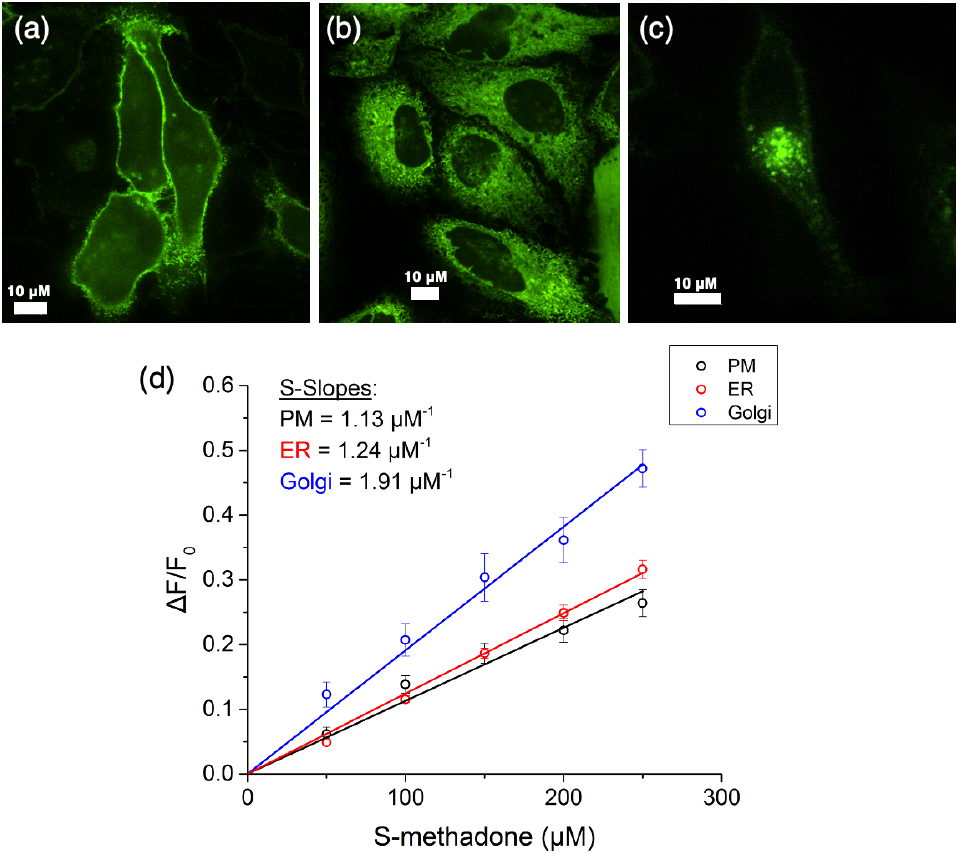
Spinning disc confocal imaging of HeLa cells transfected with (a) iS-methadoneSnFR_PM, (b) _ER, and (c) _Golgi (470 nm excitation, 535 nm emission, 100x 1.4 NA objective). Scale bar = 10 microns. (d) S-slope plotted for each organelle response at 0-250 nM S-methadone. Points are average responses to a 1 min pulse of [S-methadone]. PM n = 11 cells; ER n = 10; Golgi n = 11.

Along with other sensors of opioid signaling^11,32^, this study establishes the first genetically encoded fluorescent protein biosensor for an opioid drug, enabling real-time quantification. Furthermore, the enantioselectivity encourages biosensor development to investigate “chiral switching” of other drugs where a single enantiomer substitutes a clinically used racemate^33^. One enantiomer may serve previously unstudied indications. For example, S-methadone is now under clinical investigation as a rapidly acting antidepressant via non-opioid mechanism(s)^34^. The directed evolution results demonstrate that the nicotinic PBP may be converted to detect non-nicotinic small molecule amines by tuning residues around the aromatic 1^st^ shell. Drug biosensors *in vivo* can monitor drug concentration near receptors during administration by the experimenter or the subject, a common manipulation for studying mechanisms of reward, analgesia, and drug abuse. To meet immediate needs for diagnostics, iS-methadoneSnFR can also provide *in situ* readouts in the laboratory or home.

## Supporting information

Methadone_SI_2022.04.05

## ASSOCIATED CONTENT

### Supporting Information

The Supporting Information is available free of charge on the ACS Publications website.

Experimental details, reagents, supporting experimental figures, and amino acid and nucleotide sequences of iS-methadoneSnFR.

Constructs reported in this manuscript will be deposited in Addgene. Sequences and dose response metrics will be made available in a GitHub repository.

## AUTHOR INFORMATION

### Funding Sources

This work was supported by the National Institute on Drug Abuse (NIDA) grant DA043829, National Institute of General Medical Sciences (NIGMS) grant GM-123582, and Janelia Research Campus, HHMI. H.A.L. was supported by DA043829 and GM-123582. A.K.M was supported by DA043829, NIGMS fellowship 5T32GM007616, and National Institute of Neurological Disorders and Stroke fellowship T32NS105595. D.A.D and H.J.K were supported by the Tobacco-Related Disease Research Program (TRDRP) grant T29IR0455. A.L.N was supported by TRDRP fellowship 27FT-0022. L.L.L. and J.S.M. were supported by Janelia Research Campus, HHMI.

### Notes

A.K.M., H.A.L., L.L.L., and J.S.M. filed a patent that includes iS-methadoneSnFR.

## ACKNOWLEDGMENT

The Caltech Center for Catalysis and Chemical Synthesis supported by the Beckman Institute supported the chiral resolution work. Dr. Viviana Gradinaru and the CLOVER Center at Caltech provided plasmids and advice for AAV production. mApple-Golgi-7 was a gift from Michael Davidson (Addgene plasmid #54907). Andres Collazo and Giada Spigolon manage the Biological Imaging Facility supported by the Beckman Institute and advised on confocal imaging. The Proteome Exploration Laboratory was supported by NIH OD010788, NIH OD020013, the Betty and Gordon Moore Foundation through grant GBMF775 and the Beckman Institute at Caltech. Wei Gao, You Yu, and Heather Lukas gathered human sweat. We thank Luke Lavis for contributive discussions. We thank Aiden Aceves and Stephen Mayo for advice on docking, Theodore Chin for assistance in cloning, and Purnima Deshpande for excellent lab management.

---

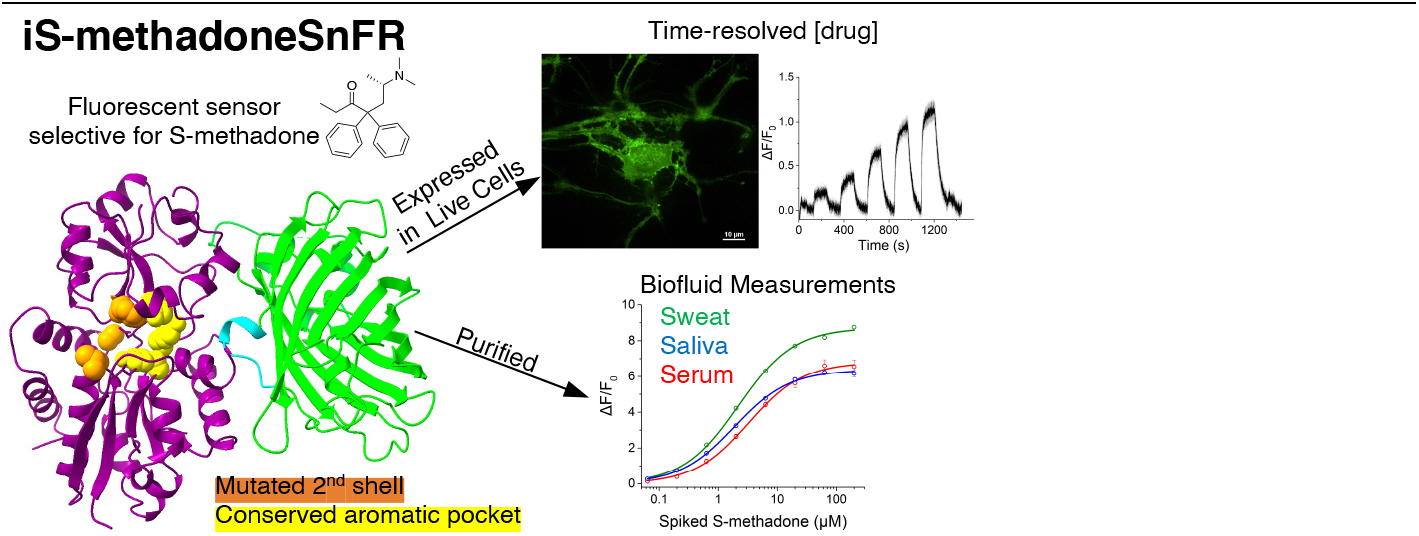

## Notes

### Competing Interest Statement

Anand Muthusamy, Henry Lester, Loren Looger, and Jonathan Marvin have filed a patent that includes iS-methadoneSnFR.

