## Supplementary material for "Three Mutations Convert the Selectivity of a Protein Sensor from Nicotinic Agonists to S-methadone for Use in Cells, Organelles, and Biofluids": Methadone_SI_2022.04.05

### Table of Contents

**I. MATERIALS, CLONING, AND ANIMAL USE STATEMENT** 2
**II. METHODS** 2
**III. SUPPORTING EXPERIMENTAL FIGURES** 7
**IV. NUCLEOTIDE AND AMINO ACID SEQUENCES OF IS-METHADONESNFR** 17
**V. REFERENCES** 18

### I. Materials, Cloning, and Animal Use Statement

Reagents

The 10x250 mm OJ-H column (Chiral Technologies, p/n 17335) was used for both analytical and preparative experiments for the chiral resolution of racemic methadone. The following reagents were purchased from Thermo Fisher Scientific: DMEM, FBS, penicillin/streptomycin solution, trypsin, DPBS, B27, Neurobasal medium, HBSS, OptiMEM, donor equine serum, and Lipofectamine 3000. The following reagents were purchased from Sigma Aldrich: ascorbic acid, BSA, racemic methadone hydrochloride, and DNase. Papain was purchased from Worthington Biochemical Corporation. Cell culture dishes were purchased from MatTek Life Sciences. HeLa and HEK293T cells were purchased from ATCC. Dpn1, Phusion polymerase, and dNTP mixture were purchased from New England Biolabs.

Cloning

We previously reported a bacterial expression vector pHHM.X513-iNicSnFR3b-(V7) (Addgene plasmid #124881; http://n2t.net/addgene:124881; RRID:Addgene_124881)[1] and mutated this plasmid during this work's directed evolution. We previously reported pMinDis.X513-iNicSnFR3a-(CC93)-ER (Addgene plasmid #125121; http://n2t.net/addgene:125121; RRID:Addgene_125121) and pMinDis.X513-iNicSnFR3a-(CC93)-PM (Addgene plasmid #125122; http://n2t.net/addgene:125122; RRID:Addgene_125122), targeting the endoplasmic reticulum and plasma membrane, respectively[1]. We cloned iS-methadoneSnFR into these mammalian expression vectors. mApple-Golgi-7 was a gift from Michael Davidson (Addgene plasmid #54907; http://n2t.net/addgene:54907; RRID: Addgene 54907) and used for the sequence targeting the fusion protein to the Golgi apparatus by appending the Golgi-targeting sequence to the biosensor's N-terminus and removing the plasma membrane targeting sequence in pMinDis.X513-iNicSnFR3a-(CC93)-PM.

Animal use statement:

C57BL/6 mice were used for a terminal cardiac puncture procedure to collect blood. Animal care was conducted in accordance with the guidelines for care and use of animals recommended by the National Institutes of Health, as stated in IACUC protocol #1386 at the California Institute of Technology. Animals were kept on a 12 h light/dark cycle and given food and water ad libitum.

### II. Methods

Chiral resolution of racemic methadone

Racemic methadone (>98% purity, Sigma Aldrich) was dissolved in ethanol containing 0.1% triethylamine and purified with ethanol containing 0.1% triethylamine on a 10x250 mm OJ-H column (Chiral Technologies, p/n 17335). From a 400 mg sample of racemic methadone hydrochloride, ~50 preparative injections afforded S-methadone, after recrystallization of each antipode from ethanol. The enantiomeric excesses of each resolved isomer were measured by supercritical fluid chromatography using eluant on 4.6x250 mm OJ-H columns. The optical rotations of the two methadone isomers agreed with published values. Optical rotation was measured in water to match each fraction to the stereochemical identity: +30.4° and -32.0° for S-methadone and R-methadone respectively compared to +26° and -26° optical rotation reference values for the *dextro-* and *levo-* isomers of the free base[2].

Docking in iNicSnFR3a

AutoDock Vina was used to perform docking[3]. The structure of iNicSnFR3a bound to varenicline was obtained from the Protein Data Bank (ID: 7S7T). The structure was prepared in AutoDockTools by removing waters, adding polar hydrogens, and assigning Gasteiger charges. Ligands were allowed torsional freedom in the docking routine. To verify the structure, varenicline was docked initially into the prepared structure. The highest scoring pose showed only a ~1 Å deviation from the nitrogens in varenicline of 7S7T. Then, S-methadone was docked into the structure. The highest scoring conformation with methadone's amine directed into the binding pocket was chosen for further analysis.

Biosensor expression by autoinduction

pHHMI plasmids bearing a biosensor gene were transformed into chemically competent BL21 (DE3) cells and grown on ampicillin plates overnight at 37 °C. Autoinduction LB was prepared according to the method of Studier 2005[4] with ampicillin (100 mg/L). A single colony was picked to inoculate each vessel with autoinduction medium. The vessel was incubated at 30 °C with shaking at 250 rpm for 28-30 h, shielded from light. Biosensor expression produced yellow-green colored cultures.

Protein purification by FPLC

Biosensors were expressed in 200 mL autoinduction cultures. Bacteria were pelleted and resuspended in 1x PBS, pH 7.4. The suspension was sonicated to lyse cells and centrifuged. The supernatant contained soluble biosensor and was applied to a Ni-NTA column on an Akta Start FPLC. The biosensor was eluted with a linear gradient from 10 to 200 mM imidazole in 1x PBS, pH 7.4. Fractions (5 mL) were collected and analyzed by SDS-PAGE to confirm purity. Pure fractions were combined and concentrated in a spin column with a 30 kDa cutoff (Amicon). The protein was buffer-exchanged into 3x PBS, pH 7.0, and concentrated to ~500 µL. Biosensor concentration was determined by absorbance at 280 nm using the extinction coefficient calculated for aromatic residues. Final pooled and concentrated protein purity was assessed by SDS-PAGE to find greater than 95% purity in samples used for both kinetic and equilibrium experiments.

Unnatural amino acid mutagenesis

"Amber codon suppression" was performed by introducing TAG codons at positions 12, 65, and 357. A permissive aminoacyl synthetase/tRNA pair (pCNF) was used to incorporate O-methyl-L-tyrosine derivatives[5]. pEVOL-pCNF and the biosensor plasmid were co-transformed into BL21 (DE3) cells. Cells were plated on double antibiotic selection plates (spectinomycin/ampicillin). A single colony was picked to inoculate a 5 mL primary LB culture, then allowed to grow overnight at 37 °C. This culture was used to inoculate a 200 mL autoinduction culture as described above. At $OD_{600}$ ~0.7, the unnatural amino acid was added to the culture dropwise while agitating. The culture was then incubated for 30-32 h with shaking at 30 °C. The biosensor protein was purified by FPLC.

Protein Digestion for Mass Spectrometry

Purified iS-methadoneSnFR samples with canonical sequence and with O-methyl-L-tyrosine substituted at positions 12 and 65 (100 µG each) was dissolved in 100 µL HEPES (50 mM, pH 8.0) containing 8 M urea. TCEP (1 µL, 500 mM in 50 mM HEPES, pH 8.0) was added, and the sample was incubated with shaking (750 rpm) at 37 °C for 20 min. 2-chloroacetamide (3 µL, 500 mM in 50 mM HEPES, pH 8.0) was added, and the sample was incubated with shaking (750 rpm) at 37 °C. Endoproteinase Lys-C (2 µL, 100 ng/µL) was added, and the sample was incubated with shaking (750 rpm) at 37 °C for 4 h. HEPES buffer (375 µL, 50 mM, pH 8.0) was added to dilute urea to a final concentration of < 2 mM. $CaCl_2$ (5 µL, 100 mM) was added, followed by trypsin (3 µL, 100 ng/µL), and the sample was incubated with shaking (750 rpm) at 37 °C overnight. The sample was acidified with TFA (15 µL, 20% v/v) and centrifuged for 30 s at 13,000 x g. Desalting was performed using ThermoFisher C18 spin columns (cat #89870) according to the manufacturer's protocol. Desalted samples were freeze-dried and stored at -20 °C prior to analysis.

Mass spectrometry validation of unnatural amino acid incorporation

Peptides were suspended in the water containing 0.2% formic acid and 2% acetonitrile for further LC-MS/MS analysis. LC-MS/MS analysis was performed with an EASY-nLC 1200 (ThermoFisher Scientific, San Jose, CA) coupled to a Q Exactive HF hybrid quadrupole-Orbitrap mass spectrometer (ThermoFisher Scientific, San Jose, CA). Peptides were separated on an Aurora UHPLC Column (25 cm × 75 µm, 1.6 µm C18, AUR2-25075C18A, IonOpticks) with a flow rate of 0.35 µL/min for a total duration of 43 min and ionized at 1.6 kV in the positive ion mode. The gradient was composed of 6% solvent B (2 min), 6-50% B (20.5 min), 50-80% B (7.5 min), 80-98% B (1 min) and 98% B (12 min); solvent A: 2% ACN and 0.2% formic acid in water; solvent B: 80% ACN and 0.2% formic acid. MS1 scans were acquired at the resolution of 60,000 from 375 to 2,000 m/z, AGC target 3e6, and maximum injection time 15 ms. The 12 most abundant ions in MS2 scans were acquired at a resolution of 30,000, AGC target 1e5, maximum injection time 60 ms, and normalized collision energy of 28. Dynamic exclusion was set to 30 s and ions with charge +1, +7, +8 and >+8 were excluded. The temperature of ion transfer tube was

275 °C and the S-lens RF level was set to 60. MS2 fragmentation spectra were searched with Proteome Discoverer SEQUEST (version 2.5, Thermo Scientific) against *in silico* tryptic digested Uniprot database of *Escherichia coli* (strain K12) and iS-methadoneSnFR1.0 protein. The maximum missed cleavages were set to 2. Dynamic modifications were set to oxidation (M, +15.995 Da), deamidation (N and Q, +0.984 Da), O-Me-Tyr (F, +30.011Da; Y, +14.016 Da), protein N-terminal acetylation (+42.011 Da) and Met-loss (-131.040 Da). Carbamidomethylation on cysteine residues (C, +57.021 Da) was set as a fixed modification. The maximum parental mass error was set to 10 ppm, and the MS2 mass tolerance was set to 0.03 Da. The false discovery threshold was set strictly to 0.01 using the Percolator Node validated by q-value. The relative abundance of parental peptides was calculated by integration of the area under the curve of the MS1 peaks using the Minora LFQ node. Spectral annotation was generated by the Interactive Peptide Spectral Annotator (IPSA, http://www.interactivepeptidespectralannotator.com/)[6] The mass spectra of the peptides containing the unnatural amino acid were taken. The fragmentation pattern and mass/charge value matched expected values in both cases.

Fluorescence dose-response relation and calculation

Biosensor and drug solutions were mixed by a liquid handling robot (epMotion) to yield 100 nM final [biosensor] and the desired [drug]. The drug plate consisted of a serial dilution of $10^{0.5}$ over each of seven steps and vehicle alone. Samples were prepared in triplicate. All solutions were 3x PBS, pH 7.0, unless otherwise stated. The plate was read using a Tecan Spark 10M with 485 nm excitation and 535 nm emission wavelengths to measure GFP fluorescence. Mean ΔF/F0 was calculated for the response to each [ligand] where $\Delta F/F_0 = (F_{drug+biosensor} - F_{biosensor})/F_{biosensor}$. Error bars are given for the standard error of the mean. The resulting data were fit with the Origin 9.2 software (OriginLabs) to the Hill equation,

$$\Delta F/\Delta F_0 = \Delta F_{max}/F_0 \left(\left[{[drug]}\big/{([drug] + EC_{50})}\right]\right)^n.$$

Because $n$ was close to 1 for measurements in solution or lysate, the S-slope,

$\Delta F/\Delta F_0 = \Delta F_{max}/F_0$ for [drug] $\ll EC_{50}$ ,

was computed as $(\Delta F_{max}/F_0)/EC_{50}$. The S-slope has dimensions $\mu M^{-1}$.

Directed evolution

Directed evolution consisted of (A) DNA library preparation, (B) culturing in 96-well plates, and (C) screening for response to ligands and obtaining winning sequences. (A) A 22-codon method was used to create mutant DNA libraries[7]. The PCR product library was transformed into TOP10 cells to amplify the DNA. Several variants from each library were sequenced to verify randomization. (B) 300 ng of the library was transformed into BL21 (DE3) cells, plated on ampicillin selection plates, and incubated overnight at 37 °C. Autoinduction medium was prepared and 800 µL were added to each well in a 96-deep well plate. A single colony was picked to inoculate each well. AeraSeal film was used to cover the plate while allowing oxygenation. The plate was incubated at 30 °C with shaking at 250 rpm for 30 h. The culture was pelleted, resuspended in 3x PBS, pH 7.0, frozen in liquid nitrogen, and thawed at room temperature to lyse bacteria. The plate was centrifuged again, providing biosensor solubilized in the lysate. (C) Lysates were transferred to a 96-well flat black plate and fluorescence in each well was read. 11 µL of 10x drug solution was added to each well and mixed by shaking. The fluorescence was measured after ligand application. $\Delta F/F_0$ was computed for each well. The top ~8 mutants were sequenced. Non-parent mutants were then transformed into BL21 (DE3) cells and used to inoculate a 10 mL autoinduction culture. The lysate was then used for a full dose response to verify the advantageous mutation.

Isothermal titration calorimetry

ITC was conducted using an Affinity ITC (TA Instruments). S-methadone stock solution and buffer-exchanged stock solution of purified biosensor were prepared using 3x PBS, pH 7.0. 40 µM of biosensor solution was added to the cell and 400 µM S-methadone (titrant) was added to the syringe. 2 µL injections of the titrant were injected at 300 s intervals 20 times. NanoAnalyze software (TA Instruments) was used to process the data. The baseline correction was applied to account for drug solvation energy. The resulting heat curve was fitted with an "independent" model to determine enthalpy, entropy, binding affinity, and stoichiometry.

Stopped-flow kinetics

Stopped-flow kinetics were measured using an Applied Photophysics SX20 stopped-flow fluorimeter with a 490 nm excitation LED and 510 nm long-pass filter at room temperature (22 ˚C). Equal volumes of 0.2 $\mu$M iS-methadoneSnFR and varying concentrations of racemic methadone were mixed (5 replicates). The first 3 ms were not analyzed to ignore mixing artifacts and instrument dead time. Data were plotted and time courses were fitted, when possible, to a single exponential approach to a plateau, using Kaleidagraph (version 4.4). $k_{obs}$ was plotted as a function of [ligand]. The linear portion of that graph was fitted, with the slope reporting $k_1$ and the y-intercept reporting $k_{-1}$. When the time course did not fit well to a single exponential component, it was fitted to the sum of two exponentials, and the faster phase ($k_{obs1}$) was treated as above to determine $k_1$ and $k_{-1}$.

Generation and analysis of racemic methadone steady-state concentration-response relation

The relaxation data were sampled at intervals of one ms. We measured the steady-state concentration-response relation for $\Delta F/F_0$ vs [racemic methadone] by taking the mean $\Delta F/F_0$ for the final 10 ms of the 1 s methadone stopped-flow relaxations. We computed $\Delta F$ by subtracting the fluorescence in methadone from that in 0 $\mu$M methadone. After correcting for instrumental offset, the value of $F_0$ was 0.05. The data were fit to the Hill equation without weighting using the nonlinear regression routine provided by the Origin 2018 software.

Analysis of R- and S-methadone interaction

To determine whether R- and S-methadone bound competitively to the sensor, we measured the effect of fixed R-methadone concentrations (0, 0.1, 1, 10, 100 $\mu$M) on the S-methadone concentration-response relation. R-methadone is not a simple competitive inhibitor because it also partially activates the sensor. Therefore, we adapted a model from enzyme kinetics for competitive inhibition with mixed alternative substrates[8]. According to this model, the following equation (1) describes fluorescence responses in the presence of both S-methadone and R-methadone,

$$\frac{\Delta F}{F_0} = \frac{Fmax_S\frac{[S]}{K_S}+Fmax_R\frac{[R]}{K_R}}{1+\frac{[S]}{K_S}+\frac{[R]}{K_R}}, \qquad \text{eq. 1}$$

where $Fmax_S$ and $Fmax_R$ are the maximum $\Delta F/F_0$ values for S- and R-methadone alone, [S] and [R] are [S-methadone] and [R-methadone], and $K_S$ and $K_R$ are the equilibrium dissociation constants for S- and R-methadone binding (in this case, $EC_{50}$s values for S- and R-methadone activation of the sensor). The data were fitted to equation 1 using the global nonlinear regression routine provided by the Origin 2018 software. The value of $F_0$ was the sensor fluorescence in buffer alone. The parameter obtained from the global fit represented the single set of $F_0$, $Fmax_S$, $Fmax_R$, $K_S$, and $K_R$ values that fitted all the data optimally.

Adeno-associated virus preparation

iS-methadoneSnFR gene was cloned into a pAAV vector with a *synapsin-1* promoter and a PDGFR plasma membrane-targeting sequence[9]. Integrity of the inverted terminal repeat sequence was confirmed by SmaI digest. Mammalian tissue culture, virus harvesting, and virus purification were performed according to the protocol of Challis 2019[10]. HEK293T cells were transfected with the pAAV, pHelper, and PHP.eB capsid genes. The medium was harvested at 3 and 5 days post-transfection and the cells were harvested at 5 days post-transfection. Digestion produced a lysate with soluble viral particles. The lysate was purified by gradient ultracentrifugation. Viral titer was determined by qPCR.

Tissue culture and transfection

HeLa cells (ATCC) were thawed and passaged twice before use in imaging studies. Cell culture followed ATCC recommended protocols. For each imaging study, 100,000 HeLa cells were plated onto a 35 mm dish with a 14 mm coverslip (MatTek) and incubated at 37 °C, 5% $CO_2$ for 24 h. Cells were then transfected with Lipofectamine 3000 using 500 ng for _PM, 250 ng for _ER, and 600 ng for _Golgi constructs in OptiMEM. Cells were kept in OptiMEM transfection medium for 24 h and then switched to standard growth medium for an additional 24 h before imaging.

Primary neuron culturing and transduction

A pregnant mouse was euthanized at embryonic day 16. The uterine sac was removed, and each embryo was decapitated before dissection. The hippocampi from several embryos were combined and digested with 15 U of papain at 37 °C for 15 min. After DNase treatment, the cells were triturated in Hanks' balanced salt solution (HBSS) with 5% donor equine serum and spun through a layer of 4% BSA and HBSS. Dishes with a 10 mm poly-D-lysine-coated glass bottom (MatTek) were coated with poly-L-ornithine and laminin 24 h prior to plating. The cells were then plated at a density of 90,000/dish in 130 µL of plating medium. After 1 h, 3 mL of complete culture medium was added to each dish. Half of the medium was changed twice a week. After 4 days, the neurons were transduced by mixing virus into the medium. After ~2 weeks, the dishes were used in imaging experiments.

Mouse serum collection

Mice were anesthetized with 5% isoflurane in air. Anesthesia was verified by slowed breathing and insensitivity to toe pinch. A needle was inserted in the left lateral thoracic wall and punctured the ventricle; 0.5-0.75 mL of blood was withdrawn into a syringe. The sample was allowed to coagulate at room temperature for 1 h and then centrifuged. The supernatant was pipetted off and used for dose response studies without any other processing.

Time-resolved measurements in cultured cells

An Olympus XI-80 microscope was equipped with an LED centered at 470 nm (LZ1- 10DB00; Led Engin), a 40-nm band-pass filter, centered at 470 nm (ET 470/40X; Chroma Technology) and an iXon DU-897 EM charge-coupled device camera (Andor Technology). Imaging was performed at 4 Hz. A programmable 8-valve perfusion system (Automate Scientific) was used to deliver solutions. Serial dilutions of S-methadone were prepared in Hanks' balanced salt solution (HBSS). PTFE-coated tubing was used to minimize gas diffusion; nonetheless, some $CO_2$ diffused out of the tubing between the solution reservoir and cell chamber, slightly alkalinizing the bicarbonate-based buffer. Hence, vehicle application without drug elicited an $F_0$ increase (Figure S8). Therefore, we used a "stuttered step" program in which each drug application was repeated to flush the solution in the tubing. The second response was taken for analysis. This response was subtracted from the mean peak response at each [S-methadone].

Analysis of cellular imaging time series data

ImageJ plugin "Time Series Analyzer" was used to calculate the average pixel intensity in the region of interest (ROI) drawn (PM, ER, or Golgi) and a background region in each frame. These data were further analyzed using the OriginLabs software. The background values were subtracted from the ROI at each frame to calculate F. A baseline was drawn with a spline to determine $F_0$ at each frame. $\Delta F/F_0$ was then calculated as $(F-F_0)/F_0$ for each frame. The steady-state response was taken as the average of the final 20 frames in each response. For the S-slope measurements, the HBSS response was subtracted from each of the responses at 50-250 nM drug. The linear fit was constrained to a y-intercept of zero.

Spinning disc confocal imaging

Images were captured using a Nikon Ti2 spinning disc confocal microscope. An environmental chamber around the stage was set to 37 °C and 5% CO2. "Perfect Focus" was used to maintain z-position before, during, and after drug solution addition. A 2x drug stock was prepared in HBSS and applied after a "baseline" image was taken. 1 min was given to allow for diffusion before capturing the post-drug image. Nikon's software was used to tile the acquisition and stitch the final image.

### III. Supporting Experimental Figures

#### Supplemental Figure 1: Chiral resolution of racemic methadone

a) Racemic methadone (peaks found at ~1.6 and ~1.8 min)

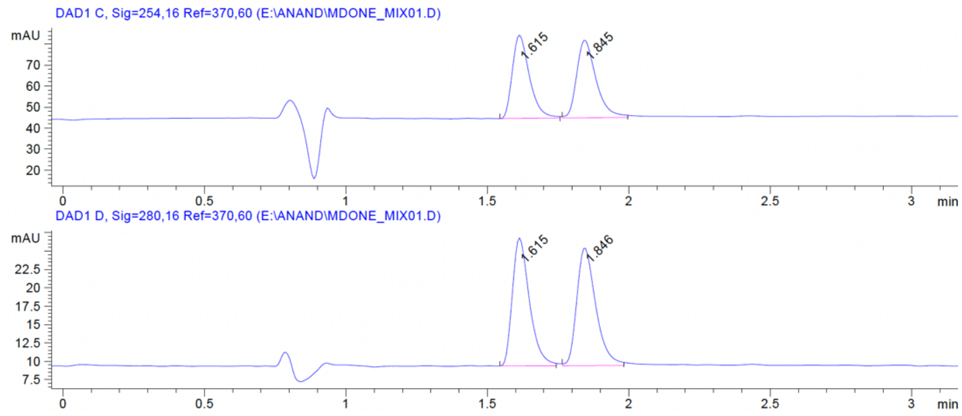


b) R-methadone (~1.6 min peak)

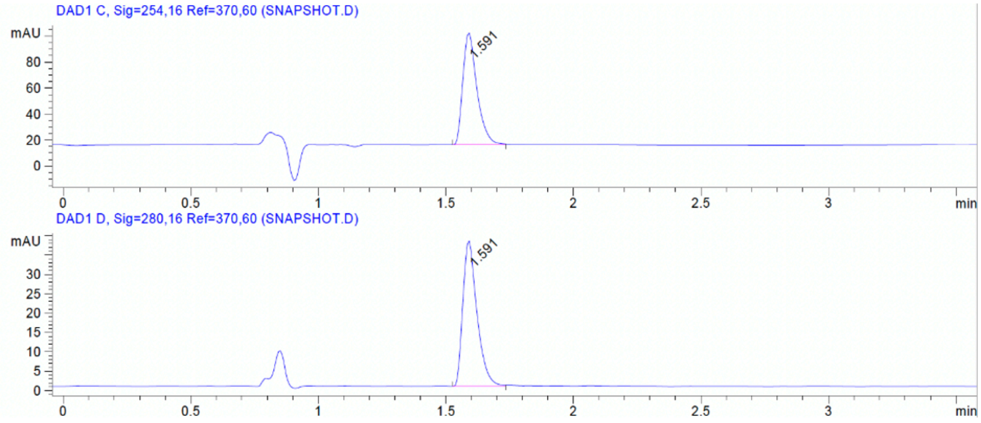


c) S-methadone (~1.8 min peak)

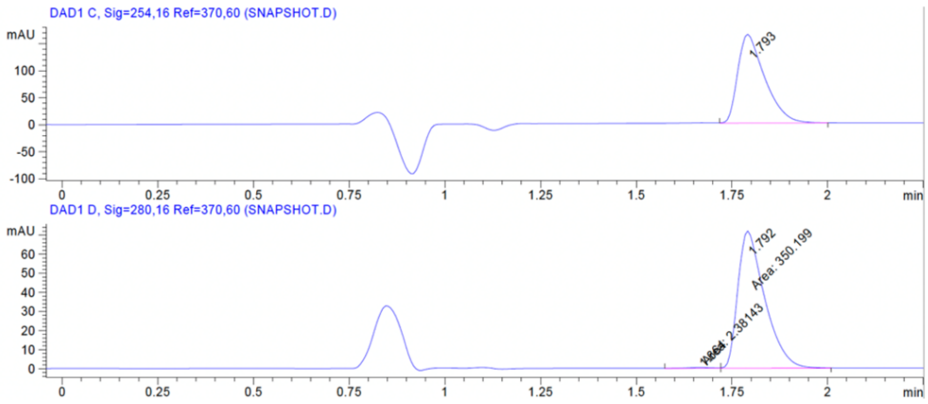


**Supplemental Figure 1**: LC-MS analysis of (a) racemic, (b) R-methadone (peak at 1.591 min) and (c) S-methadone (peak at 1.793 min) batches after chiral resolution using the same chiral column (10x250 mm OJ-H, Chiral Technologies). Each trace shows ~100% of the desired enantiomer and ~0% of the other enantiomer. Early minor peaks are from vehicle solvent. Identity of each isomer was determined by optical rotation and the sign was matched to known enantiomer assignments: +30.4° for S-methadone and -32.0° for R-methadone.

**Supplemental Figure 2: "Methadone Enantiomer x PBP Biosensor" Screen**

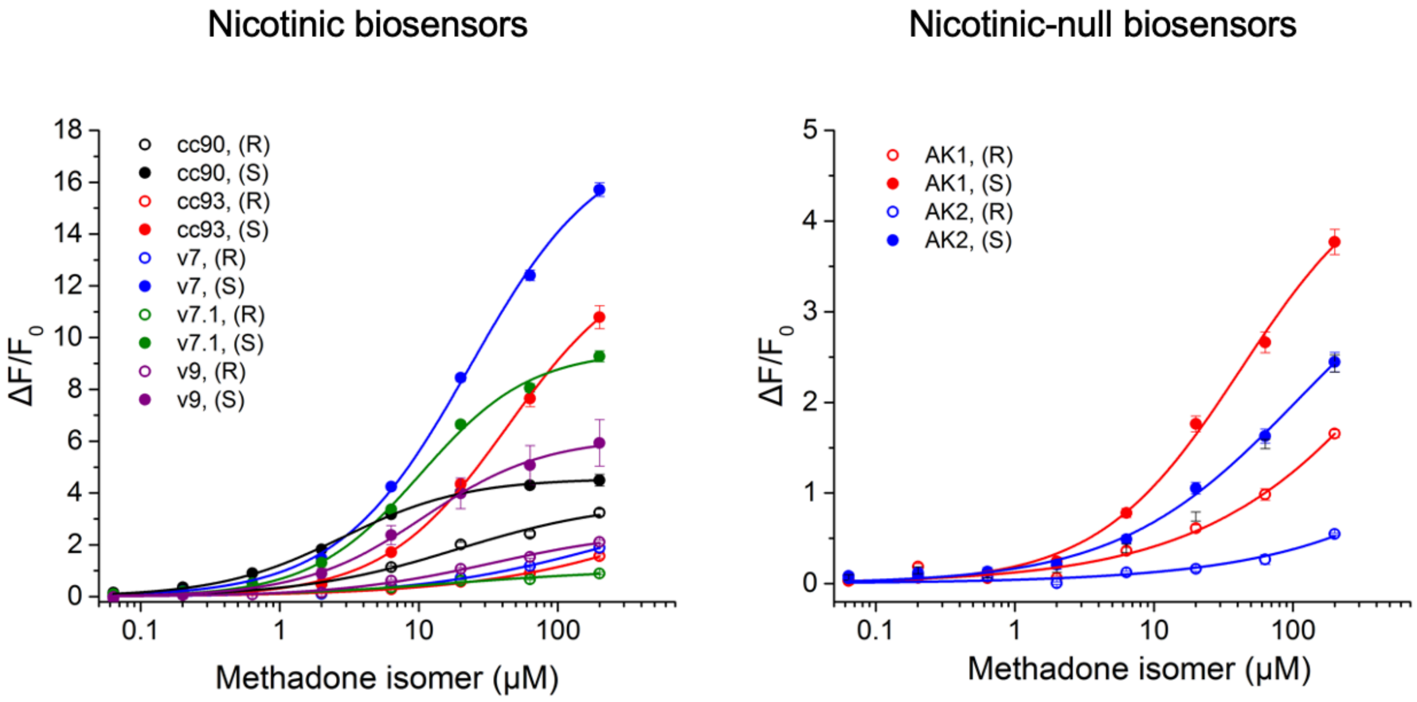


| Biosensor | R-methadone, (R) $\Delta F_{max}/F_0$ | $EC_{50}$ | $n_H$ | S-Slope | S-methadone, (S) $\Delta F_{max}/F_0$ | $EC_{50}$ | $n_H$ | S-slope |
|---|---|---|---|---|---|---|---|---|
| cc90 = iNicSnFR2 | 3.7 | 19.2 | 0.77 | 0.2 | 4.6 | 2.8 | 0.98 | 1.6 |
| cc93 = iNicSnFR3a | - | - | - | <0.1 | 13.2 | 43.9 | 0.97 | 0.3 |
| v7 = iNicSnFR3b | 3.8 | 211.8 | 0.64 | <0.1 | 17.8 | 23.3 | 0.92 | 0.8 |
| v7 43V | 1.1 | 18.4 | 0.65 | <0.1 | 9.5 | 10.3 | 1.10 | 0.9 |
| v9 = iAChSnFR | 2.6 | 33.2 | 0.79 | <0.1 | 6.1 | 10.9 | 1.00 | 0.6 |
| AK1 | - | - | - | <0.1 | 4.7 | 42.2 | 0.85 | 0.1 |
| AK2 = iSketSnFR2 | - | - | - | <0.1 | 4.1 | 110 | 0.67 | <0.1 |

**Supplemental Figure 2**: Dose-response relations for previously developed biosensor variants against R-methadone and S-methadone. Top left: nicotinic biosensors originally developed for iNicSnFR and iAChSnFR campaigns[1,11]. Top right: nicotinic-null biosensors originally developed for iSKetSnFR (ketamine biosensor) campaigns[12]. Table: Hill fit parameters for every biosensor-methadone enantiomer pair dose response. iNicSnFR3b displayed the greatest S-slope for both S-methadone and R-methadone while preserving dynamic range of $\Delta F_{max}/F_0 > 10$.

**Supplemental Figure 3: Evolution Tree: iNicSnFR3a to iS-methadoneSnFR**

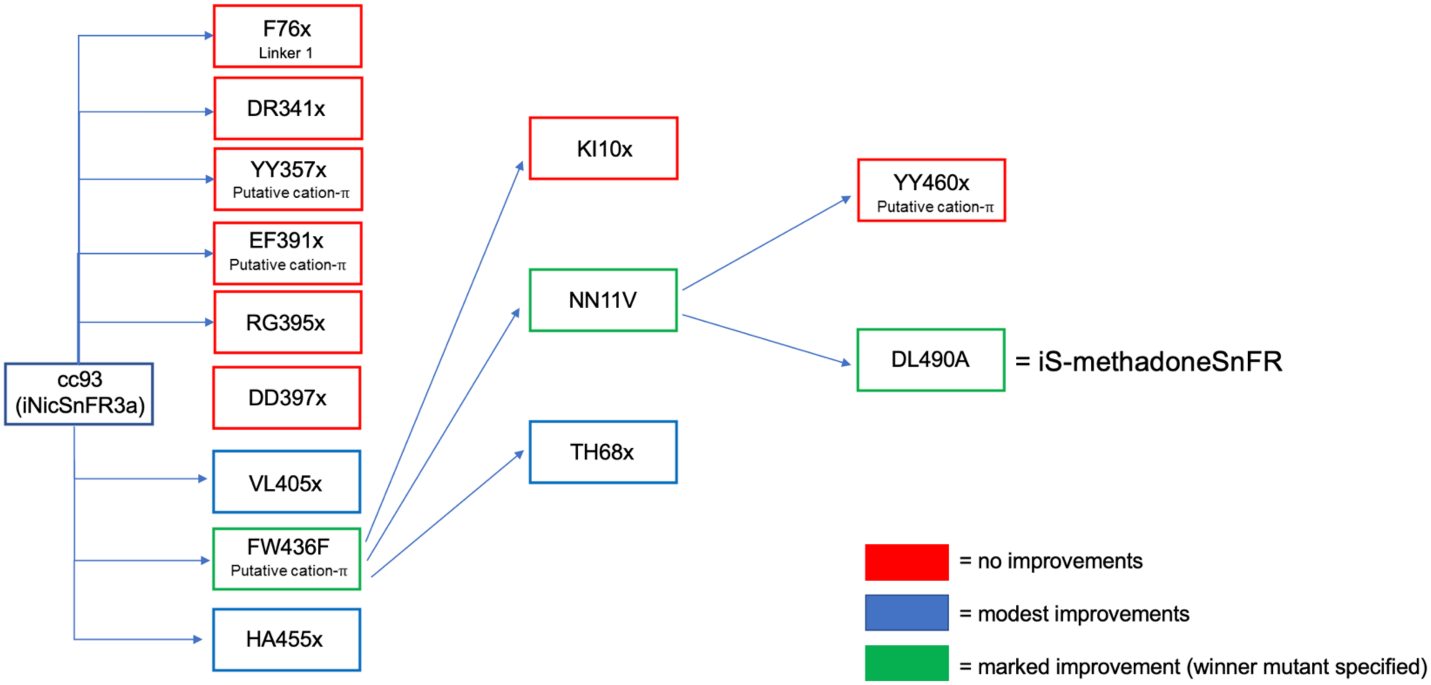


**Supplemental Figure 3:** Evolution tree from iNicSnFR3a to iS-methadoneSnFR. Residue nomenclature: first and second residues before the position number are the amino acids in the OpuBC homologue from *Thermoanaerobacter sp* X513 and iNicSnFR3b, respectively. Functional role of the residue is noted. Each arrow and box pair represents a single residue site-saturation experiment. Red outlined boxes indicate positions that yielded only variants inferior to the parent. Blue outlined boxes indicate residues that yielded variants with modest improvements, typically 10-20% increases in S-slope. Green outlined boxes indicate mutations that yielded marked improvements accepted for additional mutagenesis rounds or, finally, for iS-methadoneSnFR.

**Supplemental Figure 4: Unnatural amino acid substitution at positions 12 and 65**

(a)

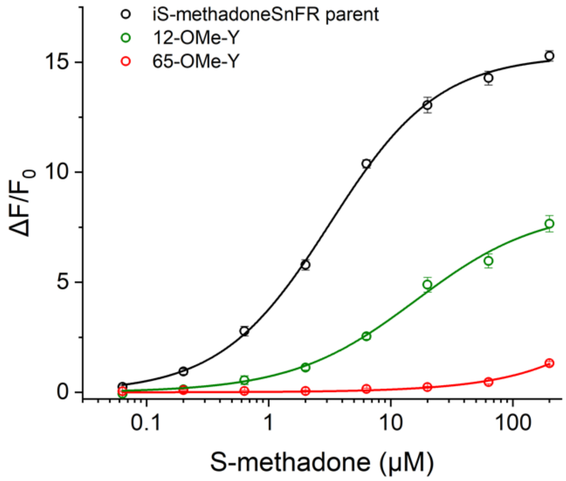


(b)

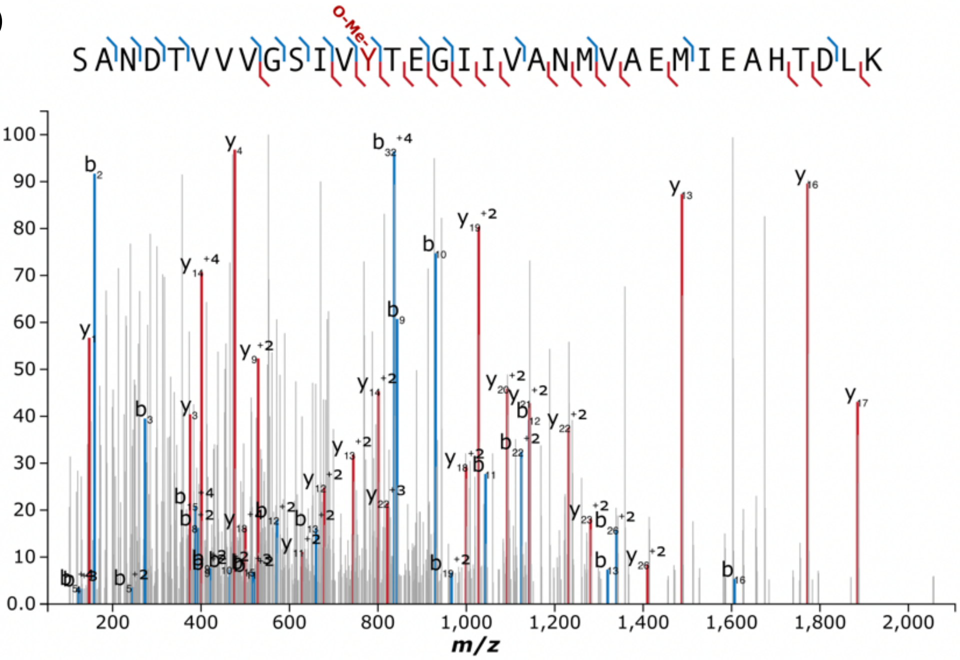


| UAA Fragment | Charge (z) | Experimental m/z | Theoretical m/z | Mass Error (Da) |
|---|---|---|---|---|
| b13 | 1 | 1319.6825 | 1319.6845 | -0.002 |
| b13 | 2 | 660.3418 | 660.3459 | -0.0041 |
| b15 | 3 | 517.2863 | 517.2631 | 0.0232 |
| b15 | 4 | 388.1996 | 388.1992 | 0.0004 |
| b16 | 1 | 1606.8046 | 1606.7962 | 0.0083 |
| b19 | 2 | 966.521 | 966.52 | 0.001 |
| b22 | 2 | 1124.5962 | 1124.5803 | 0.0159 |
| b26 | 2 | 1339.6737 | 1339.6746 | -0.0009 |
| b32 | 4 | 836.9033 | 836.9153 | -0.012 |
| y22 | 2 | 1231.6143 | 1231.6191 | -0.0048 |
| y22 | 3 | 821.3989 | 821.4151 | -0.0163 |
| y23 | 2 | 1281.1467 | 1281.1533 | -0.0066 |
| y26 | 2 | 1409.694 | 1409.7221 | -0.0281 |

(c)

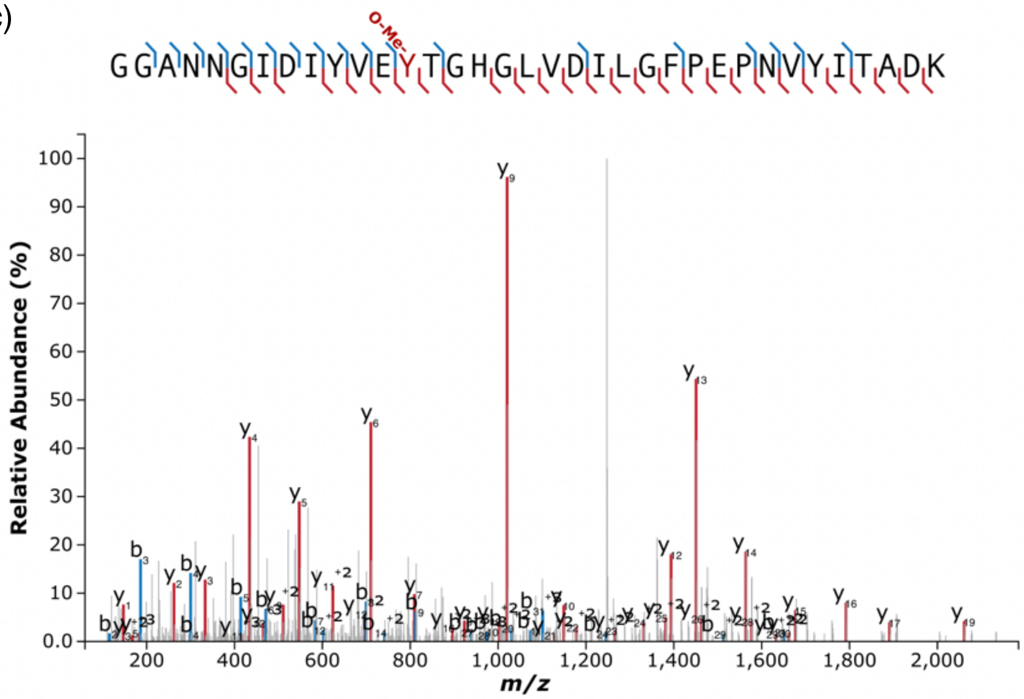


| UAA Fragment | Charge (z) | Experimental m/z | Theoretical m/z | Mass Error (Da) |
|---|---|---|---|---|
| b20 | 2 | 1030.4974 | 1030.4898 | 0.0077 |
| b27 | 3 | 938.4449 | 938.4643 | -0.0194 |
| b31 | 2 | 1651.8505 | 1651.8222 | 0.0282 |
| b31 | 3 | 1101.5284 | 1101.5506 | -0.0221 |
| y26 | 2 | 1462.7508 | 1462.7396 | 0.0112 |
| y28 | 2 | 1576.8031 | 1576.7951 | 0.008 |

**Supplemental Figure 4:** Amber suppression unnatural amino acid mutagenesis was used to incorporate the unnatural amino acid (UAA) O-methyltyrosine into positions 12 and 65 in separate constructs. (a) The dose responses for these two mutants are compared to the fully canonical sequence. Substitution at position 12 decreases dynamic range but roughly maintains $EC_{50}$; however, methylation of 65Y sidechain led to a near-null mutant. (b) Mass spectrometry validation of UAA incorporation at position 12: peptide containing the UAA was identified by mass fragments. 'b' fragments (blue) are numbered for the fragment length starting from N-terminal end of the peptide. 'y' fragments (red) are numbered for the fragment length starting from the C-terminal end of the peptide. The table lists the expected and theoretical mass difference for peptide fragments containing the UAA. (c) Mass spectrometry validation of UAA incorporation at position 65 using the same method in (b).

**Supplemental Figure 5: Analysis of dose-response relations for R-methadone and S-methadone mixtures**

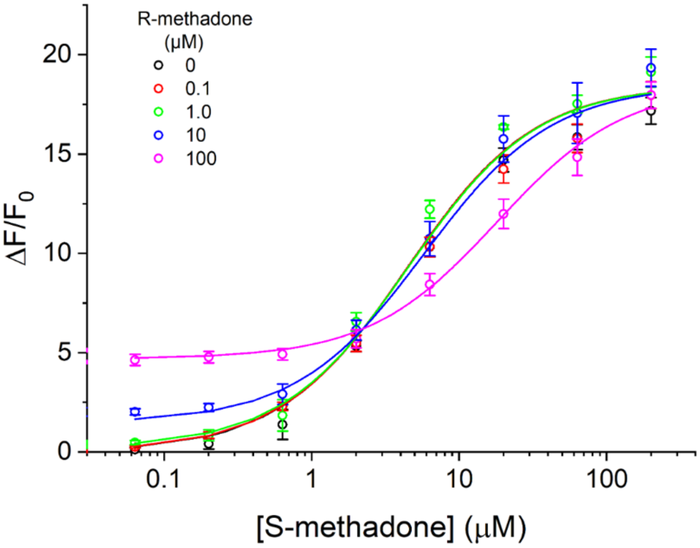


$$\frac{\Delta F}{F_0} = \frac{Fmax_S \frac{[S]}{K_s} + Fmax_R \frac{[R]}{K_R}}{1 + \frac{[S]}{K_S} + \frac{[R]}{K_R}},$$

| [R-methadone] (μM) | Fit $R^2$ |
|---|---|
| 0 | 0.987 |
| 0.1 | 0.992 |
| 1.0 | 0.989 |
| 10 | 0.991 |
| 100 | 0.995 |

**Supplemental figure 5: Interactions between R- and S-methadone at iS-methadoneSnFR.** Dose-response relations were measured for S-methadone in the additional presence of five R-methadone concentrations. A model of competitive inhibition with mixed alternative substrates was used to account for R-methadone and S-methadone competitive binding with partial R-methadone agonism. Consistent with the model, R-methadone right-shifted the [S-methadone] concentration-response relation but did not significantly affect the maximum response.

**Supplemental Figure 6: Stopped-flow – raw data and 1 sec steady-state fit**

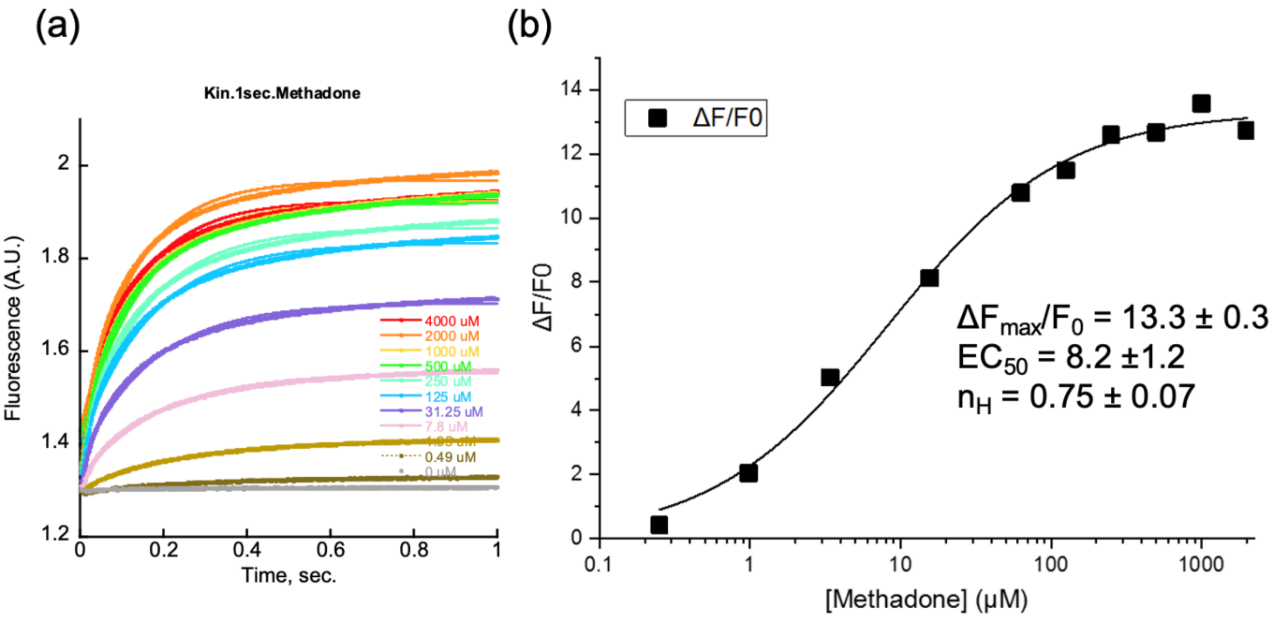


**Supplemental Figure 6:** (a) 1 s stopped-flow data. Racemic methadone was mixed with purified iS-methadoneSnFR in a chamber while monitoring fluorescence. Concentrations listed are twice the final [S-methadone]. (b) The mean response for the final 10 ms of the relaxation [methadone] was fitted to the Hill equation. The $EC_{50}$ of 8.2 µM is ~ double that of the fluorescence dose-response $EC_{50}$ measured for S-methadone alone (Fig. 4B).

**Supplemental Figure 7: pH Dependence of iS-methadoneSnFR dose-response**

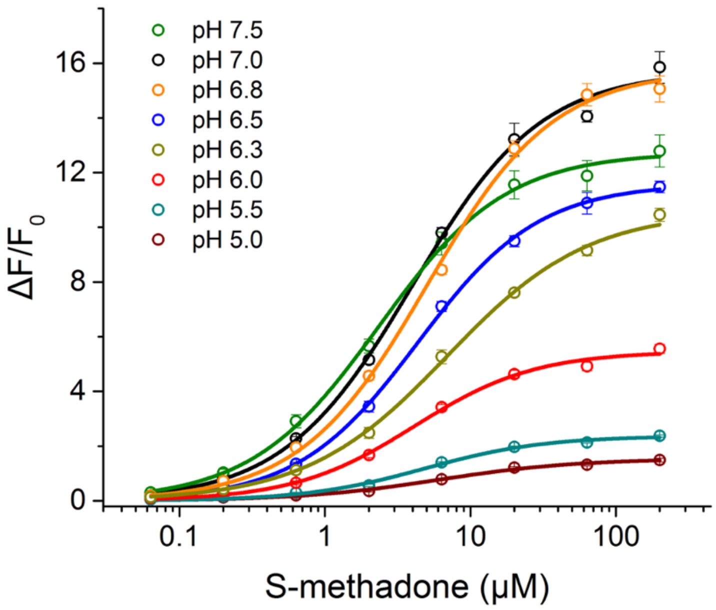


| pH | $\Delta F_{max}/F_0$ | $EC_{50}$ | $n_H$ | S-Slope |
|---|---|---|---|---|
| 7.5 | 12.7 | 2.33 | 0.99 | 5.45 |
| 7.0 | 15.7 | 3.97 | 0.98 | 3.95 |
| 6.8 | 15.8 | 5.03 | 1.00 | 3.14 |
| 6.5 | 11.6 | 4.36 | 1.05 | 2.66 |
| 6.3 | 10.6 | 6.92 | 0.9 | 1.53 |
| 6.0 | 5.5 | 4.12 | 1.05 | 1.33 |
| 5.5 | 2.4 | 4.85 | 1.06 | 0.49 |
| 5.0 | 1.5 | 6.03 | 0.9 | 0.25 |

**Supplemental Figure 7**: Effect of pH on the S-methadone dose-response relation. 3x PBS buffers were prepared from pH 5.0 to 7.5 in half-unit increments. Dose-response data were collected in each buffer and plotted. Hill fit parameters and computed S-slope are given in the righthand table. Like other GFP-based biosensors, the iS-methadoneSnFR response decreased at acidic pH. The S-slope remained > 1 $\mu M^{-1}$ at pH 6.0, enabling measurements across the Golgi pH range. S-slope at pH 7.5 was 1.7x larger than that at pH 6.8 and 2.6x larger than that at pH 6.3.

**Supplemental Figure 8: HeLa organelle study – bath perfusion dose-response**

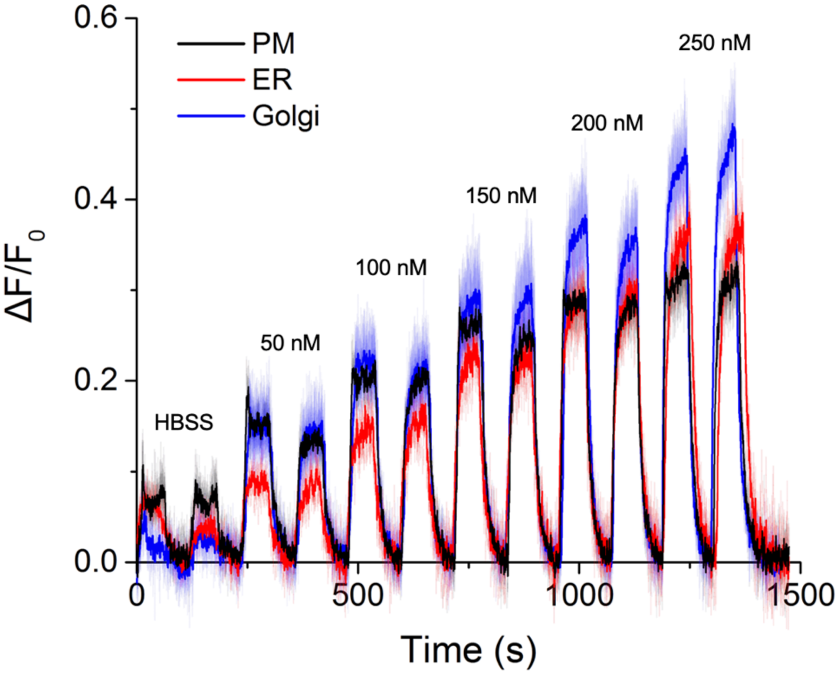


**Supplemental Figure 8**: Time-resolved dose-response data from imaging experiments for iSmethadoneSnFR targeted to various HeLa cellular compartments. A “stuttered step” perfusion method (1 min S-methadone on, 1 min wash, each dose applied twice) was used to minimize pH effects. Fluorescence images (40x, 1.0 NA, 470 nm excitation) were acquired at 4 Hz. Traces show mean responses (PM n = 11 cells; ER n = 10; Golgi n = 11). Data were smoothed using a 4-point moving average. SEM denoted by faint bands.

**Supplemental Figure 9: Validation in primary hippocampal neurons**

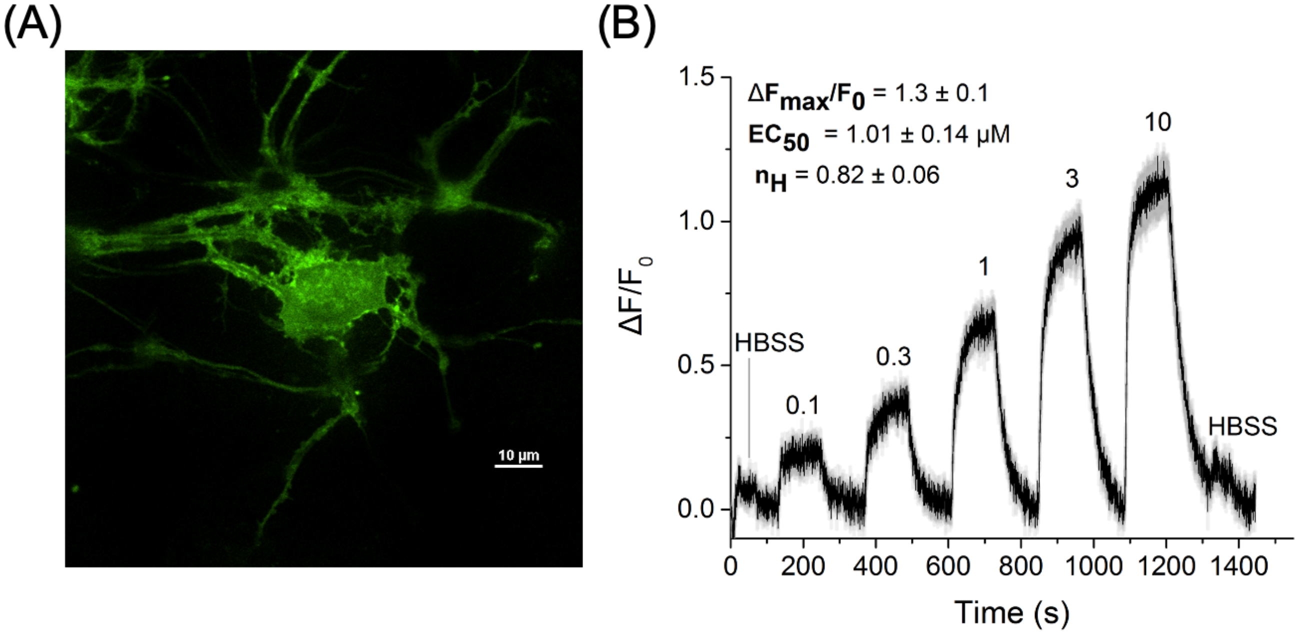


**Supplemental Figure 9**: (a) Spinning disc confocal imaging of a cultured mouse hippocampal neuron transduced with PHP.eB-*hSyn*-iS-methadoneSnFR-PM-WPRE (100x, 1.4 NA objective; 488 nm excitation, 535 nm emission. Scale bar = 10 µm.) (b) Responses to pulses of S-methadone (2 min drug application followed by 2 min rinse). iS-methadoneSnFR detected S-methadone in neuronal cultures across the pharmacologically relevant range (50 nM to 3 µM). Traces are mean response ± SEM (n = 12 neurons, SEM as gray bounds). S-methadone concentration is given above traces (in µM). The final 10 s of the S-methadone response was averaged across the cells and the response to vehicle alone (HBSS) was subtracted to measure the dose-response relation. The Hill fit parameters of the dose-response were $\Delta F_{max}/F_0$ = 1.3 ± 0.1, $EC_{50}$ = 1.01 ± 0.14 µM, and $n_H$ = 0.82 ± 0.06.

### IV. Nucleotide and amino acid sequences of iS-methadoneSnFR

iS-methadoneSnFR nucleotide sequence:

ATGCATCATCATCATCATCATGGTTATCCCTATGATGTTCCAGATTATGCTGGGGCCCAGCCGGCCAGATC
TGCGAACGACACCGTAGTTGTGGGCTCGATCGTGTTTACAGAAGGGATTATCGTCGCAAACATGGTGGCA
GAGATGATTGAGGCGCATACAGACCTTAAGGTGGTTCGCAAACTGAACCTTGGCGGGGAGAACGTTAACT
TTGAAGCCATTAAACGCGGAGGTGCGAATAATGGTATTGACATTTACGTGGAGTACACTGGGCACGGTCT
TGTGGATATTCTGGGGTTCCCGGAGCCGAACGTCTATATCACCGCCGACAAGCAGAAGAACGGCATCAA
GGCGAACTTCAAGATCCGCCACAACGTGGAGGACGGCAGCGTGCAGCTCGCCGACCACTACCAGCAGA
ACACCCCCATCGGCGACGGCCCCGTGCTGCTGCCCGACAACCACTACCTGAGCACCCAGTCCGTGCTGA
GCAAAGACCCCAACGAGAAGCGCGATCACATGGTCCTGCTGGAGTTCGTGACCGCCGCCGGGATCACTC
TCGGCATGGACGAGCTGTACAAGGGCGGTACCGGAGGGAGCATGAGCAAGGGCGAGGAGCTGTTCACC
GGGGTGGTGCCCATCCTGGTCGAGCTGGACGGCGACGTAAACGGCCACAAGTTCAGCGTGCGCGGCGA
GGGCGAGGGCGATGCCACCAACGGCAAGCTGACCCTGAAGTTCATCTGCACCACCGGCAAGCTGCCCG
TGCCCTGGCCCACCCTCGTGACCACCCTGACCTACGGCGTGCAGTGCTTCAGCCGCTACCCCGACCACA
TGAAGCAGCACGACTTCTTCAAGTCCGCCATGCCCGAAGGCTACGTCCAGGAGCGCACCATCAGCTTCA
AGGACGACGGCACCTACAAGACCCGCGCCGAGGTGAAGTTCGAGGGCGACACCCTGGTGAACCGCATC
GAGCTGAAGGGCATCGACTTCAAGGAGGACGGCAACATCCTGGGGCACAAGCTGGAGTACAACTTTCCG
CCGCCCAGCTCTACTGATCCAGAAGGTGCATACGAAACCGTGAAGAAGGAGTACAAACGTAAATGGAATA
TTGTATGGCTCAAACCACTGGGATTCAACAATACGTATACGCTTACCGTTAAAGACGAACTGGCGAAACAG
TATAACCTTAAAACCTTCAGTGACTTAGCGAAAATCTCGGATAAGCTGATTCTGGGTGCAACGATGTTCTTT
TTAGAAGGGCCCGATGGTTACCCAGGCCTGCAAAAACTGTACAATTTCAAATTCAAGCACACCAAAAGCAT
GGACATGGGTATTCGCTATACCGCCATTGATAATAACGAAGTTCAGGTAATTGATGCCTTCGCCACTGATG
GCTTGCTGGTGAGCCACAAATTAAAAATTCTGGAGGATGATAAAGCGTTCTTCCCGCCGTATTATGCTGCC
CCCATCATCCGTCAGGATGTCTTAGATAAGCATCCTGAACTGAAGGACGTGCTGAACAAACTCGCGAATC
AAATTTCAGCGGAAGAAATGCAGAAACTGAATTACAAGGTGGACGGTGAGGGTCAGGACCCAGCGAAAG
TAGCTAAGGAGTTTTTGAAAGAGAAAGGTTTAATTCTGCAGGTCGACGAACAAAAACTCATCTCAGAAGAG
GATCTGAATTAA

iS-methadoneSnFR amino acid sequence:

His6-tag
Myc tag
PBP
Linkers
cpGFP

MHHHHHHGYPYDVPDYAGAQPARSANDTVVVGSIVFTEGIIVANMVAEMIEAHTDLKVVRKLNLGGENVNFEAI
KRGGANNGIDIYVEYTGHGLVDILGFPEPNVYITADKQKNGIKANFKIRHNVEDGSVQLADHYQQNTPIGDGPV
LLPDNHYLSTQSVLSKDPNEKRDHMVLLEFVTAAGITLGMDELYKGGTGGSMSKGEELFTGVVPILVELDGDV
NGHKFSVRGEGEGDATNGKLTLKFICTTGKLPVPWPTLVTTLTYGVQCFSRYPDHMKQHDFFKSAMPEGYVQ
ERTISFKDDGTYKTRAEVKFEGDTLVNRIELKGIDFKEDGNILGHKLEYNFPPPSSTDPEGAYETVKKEYKRKW
NIVWLKPLGFNNTYTLTVKDELAKQYNLKTFSDLAKISDKLILGATMFFLEGPDGYPGLQKLYNFKFKHTKSMD
MGIRYTAIDNNEVQVIDAFATDGLLVSHKLKILEDDKAFFPPYYAAPIIRQDVLDKHPELKDVLNKLANQISAEEM
QKLNYKVDGEGQDPAKVAKEFLKEKGLILQVDEQKLISEEDLN
